# And the best task is …? Using Task potency to infer task specificity

**DOI:** 10.1101/111187

**Authors:** Roselyne J. Chauvin, Maarten Mennes, Alberto Llera, Jan K. Buitelaar, Christian F. Beckmann

## Abstract

When an individual engages in a task, the associated evoked activities build upon already ongoing activity, itself shaped by an underlying functional connectivity baseline (Fox et al., 2009; Smith et al., 2009; Tavor et al., 2016). To facilitate understanding the building blocks of cognition we incorporate the idea that task-induced functional connectivity modulation with respect to its underlying resting state functional connectivity is task-specific. Here, we introduce a framework incorporating *task potency*, providing direct access to task-specificity through enabling direct comparison between task paradigms. In particular, to study functional connectivity modulations related to cognitive involvement in a task we define task potency as the amplitude of connectivity modulations away from the brain’s baseline functional connectivity architecture as observed during a resting state acquisition. We demonstrate the use of our framework by comparing three tasks (visuo-spatial working memory, reward processing, and stop signal task) available within a large cohort. Using task potency, we demonstrate that cognitive operations are supported by a common baseline of within-network interactions, supplemented by connections between large-scale networks in order to solve a specific task.

**Highlights:**

- Task potency framework defines modulation of functional connectivity away from baseline resting state
- More within-than between-network modulations are induced by task performance
- Between-network modulations are task-specific
- Edges modulated by multiple tasks are mostly within-network
- The task potency can be used to define the most potent task

## 1. Introduction

Advances in functional brain imaging have provided tremendous insight into the neural correlates of cognition by relating behavioural descriptions to local changes in brain activity via oxygen metabolism using functional Magnetic Resonance Imaging (fMRI). Typical experimental studies probe specific cognitive functions (Geyer et al., 2011; Zilles and Amunts, 2010), and thereby inform about the *sensitivity* of brain areas to the experimental manipulation of interest (Aguirre et al., 2002; Harley, 2004). Yet, single neuroimaging studies do not allow making inferences about whether an observed area exclusively responds to cognitive function A or whether it is also sensitive to manipulation of function B. As such, they cannot inform about the *specificity* of a brain area for the tested cognitive function.

To be informative about specificity rather than mere sensitivity and thus allow for reverse inference (Aguirre et al., 2002; Poldrack, 2006), study participants would need to be probed for various cognitive functions across a broad repertoire of domains. Such multi-paradigm investigations are technically and logistically challenging and therefore remain rare. Accordingly, in order to indirectly infer specific behavioural relevance for the neural responses they observed, authors typically resort to previously reported results via literature meta-analysis or alternative initiatives that validate study results (Hutzler, 2014; Poldrack, 2011; Schwartz et al., 2013; Varoquaux and Thirion, 2014). Yet, such literature-based techniques are troubled by typical biases associated with the publication process including article imprecision, ‘File Drawer’ issues, and p-value tweaking (Button et al., 2013; Poldrack, 2006).

In all of these approaches, the collection of alternative tasks directly measured or inferred upon through meta-analysis provide a *functional baseline* for comparison that gives rise to the notion of task specificity in the sense that effects size estimates are elevated relative to this basal state. Corroborating the idea of such a functional baseline we here propose to take advantage of the fact that task-related activity patterns can be delineated purely from resting state functional MRI (rFMRI) data (Smith et al., 2009) in that rFMRI data exhibits dynamics that correspond to the major functional activation patterns that can be derived across a vast repertoire of tasks. Such baseline network structure supports the idea that specific cognitive states are produced by a baseline of common network activity supplemented by specific modulations across the brain’s functional architecture, rather than being orchestrated by independent activity in single regions. An important corollary is that cognitive function emerges by embedding unique regional activity within the context of larger network processes, as described in the ‘massive redeployment hypothesis’ (Anderson, 2007; Lloyd, 2000). Hence, we propose to utilise rFMRI-derived networks as a surrogate of a broad set of different tasks probing different cognitive domains that allow to formulate an innovative framework for disentangling sensitivity and specificity.

Our approach focuses on task *sensitivity*, task s*pecificity*, and task *potency* at the level of mesoscopic functional connectivity within the brain. Functional connectivity can successfully differentiate between mental states (Shirer et al., 2012), corroborating the idea that mental states can be defined based on specific connectivity profiles, similar to a “brain fingerprint” (Cole et al., 2016; Yeo et al., 2011). Accordingly, even more than local brain activity, distributed patterns of brain activity might be the best reflection of mental states and behavioural performance. As an example, a region’s task involvement is related to its whole brain connectivity pattern, where a region’s connectivity strength with the task-positive network effectively determines its task-related activity (Mennes et al., 2010). In another example, memory-learning performance could be accurately predicted based on whole-brain connectivity, where connections between regions not involved in the task were equally important predictors (Yamashita et al., 2015). This evidence suggests that both general as well as task-specific neural processes support task performance.

We describe this framework in the context of emerging large cohort functional imaging studies that involve multiple experimental fMRI designs along with rFMRI measures, allowing for within-subject comparisons between cognitive paradigms relative to the resting condition. Examples include the Human Connectome Project (Glasser et al., 2016; Van Essen et al., 2012) aimed at deciphering the complex relationship between brain functions, cognition, and the functional and structural human connectome within a normal cohort. Similar projects are translating these efforts to the clinical domain (e.g., NeuroIMAGE (Rhein et al., 2015), PNC (Satterthwaite et al., 2016), ABCD (Bjork et al., 2017)). Through directly comparing estimates from tasks relative to rest we capitalize on the increased statistical sensitivity to effect size differences that within-subject designs offer and introduce the concept of *task potency*, which allows for characterisations of differential effect size in relation to different types of experimental manipulation. We utilise this to disentangle general versus task-specific processes by indexing the presence or absence of significant functional connectivity under different tasks and posit that generic neural processes occur across multiple tasks and will be small in differential potency. Specific neural processes, however, will be related to a single cognitive process and have large associated differential potency. Of note, under this framework a specific cognitive process could be of importance for solving several tasks.

In this manuscript we considered three tasks, yet the framework is readily extensible to other tasks. We linked connectivity changes to regional task specialisation and involvement at the network level by defining functional connectivity fingerprints using a top-down, functionally defined, hierarchical atlas that divides 11 larger resting-state networks into 184 smaller regions using Instantaneous Connectivity Parcellation (ICP, (van Oort et al., 2017)). After defining edges, i.e., connections, that are relevant for a task, our approach allowed differentiating experimental designs in their specificity of modulating these connections (i.e., is this change in brain connectivity specifically associated with task A or task B?). Using effect-sizes on population distributions of these connectivity deviations, we quantified the potency of each task for selected functional connections. As such, the amplitude of modulation becomes a feature that defines task specificity as a new tool to do reverse inference and to understand the relative effect of one task within a larger battery of tasks.

## 2. Methods

### 2.1. Participants

We use MRI data from the NeuroIMAGE sample (N total > 800 participants; see von Rhein et al., 2014). In the current analyses we included data from healthy control participants only (initial N=385) who each performed at least one of the following tasks during fMRI scanning: response inhibition (Stop Signal Task (STOP), (Logan et al., 1984; Rhein et al., 2015; van Rooij et al., 2015)), reward processing (REWARD, (Hoogman et al., 2011; Knutson et al., 2001; Rhein et al., 2015; von Rhein et al., 2015)), spatial working memory (WM, (van Ewijk et al., 2015; Klingberg et al., 2002; McNab et al., 2008; Rhein et al., 2015)) (see supplementary table 1). In addition, to the task-based MRI scans each participant completed a task-free resting state fMRI sca. All participants also completed an anatomical scan for registration purposes. MRI acquisition parameters are shown in table 1.

Functional scans exhibiting limited brain coverage or excessive head motion were excluded from further processing. Limited brain coverage was defined as having less than 97% overlap with the MNI152 standard brain after image registration. Applying this criterion excluded 47 subjects (details in table 1). In addition, we excluded from each task those participants who were among top 5% in terms of head motion as quantified by RMS-FD, the root mean square of the frame-wise displacement computed using MCFLIRT (Jenkinson et al., 2002). After applying these criteria, we selected only participants that completed at least a task and a resting state scan resulting in the inclusion of data from 218 healthy controls, comprising 218 resting state acquisitions, 111 STOP acquisitions, 123 REWARD acquisitions, and 147 WM acquisitions. Participants ranged in age between 8.6 and 30.5; mean=17.0; sd=3.5; 45.9% were male.

**Table 1:** MRI acquisition parameters and participant characteristics (demographic distributions are represented in supplementary figure 1)

|  |  |  |  |  |
| --- | --- | --- | --- | --- |
| **T1-weighted structural MRI parameters** | T1-weighted MPRAGE, TR=2730 ms, TE=2.95 ms, T1=1000ms, flip angle=7, matrix size=256x256, FOV=256mm, 176 slices with 1mm isotropic voxels |  |  |  |
| **fMRI image acquisition parameters** |  |  |  |  |
|  | **rFMRI** | **STOP** | **REWARD** | **WM** |
| General parameters | TE=40 ms, FOV=224mm, 37 axial slices, flip angle=80, matrix size=64x64, in-plane resolution=3.5mm, slice thickness/gap=3.0mm/0.5mm |  |  |  |
| N volumes | >260 | 86 * 4 blocks | >300 | 107 * 4 blocks |
| TR in ms<sup>1</sup> | 1960 ( | 2340 | 2340 | 2340 |
| N first volumes rejected<sup>2</sup> | 5 | 4 | 5 | 3 |
| **Participant characteristics** |  |  |  |  |
| N initial | 302 | 239 | 256 | 266 |
| N rejected for limited brain coverage | 73 | 4 | 8 | 6 |
| N rejected for head motion | 11 | 11 | 12 | 13 |
| N who also completed a resting state scan and that were used in final analyses | 218 | 111 | 123 | 144 |
| RMS-FD min-max | 0.026 - 1.930 | 0.025 - 0.947 | 0.027 - 0.976 | 0.029 - 1.504 |
| RMS-FD mean (std) | 0.171 (0.224) | 0.093 (0.098) | 0.145 (0.145) | 0.149 (0.191) |
| Age min - max | 8.6 - 30.5 | 8.6 - 27 | 9.1 - 30.1 | 8.6 - 30.1 |
| Age mean (std) | 17 (3.5) | 17.4 (3.6) | 17.6 (3.7) | 17.4 (3.6) |
| % male | 45.8% | 45.5% | 44.5% | 47% |

<sup>1</sup> Except 1 scan which was done at 1860ms in rFMRI, 1 at 2150ms in STOP, and 1 at 2280ms in REWARD
<sup>2</sup> The number of initial volumes removed from further analyses varied to ensure comparability with earlier studies that used these data. This variation will have very limited impact on the current analyses.

### 2.2. fMRI Preprocessing

All fMRI acquisitions were processed using tools from FSL 5.0.6. (FSL; <u>http://www.fmrib.ox.ac.uk/fsl</u>; (Beckmann and Smith, 2004; Jenkinson et al., 2012; Woolrich et al., 2009)). We employed the following pipeline: removal of the first volumes to allow magnetization equilibration (see table 1), head movement correction by volume-realignment to the middle volume using MCFLIRT, global 4D mean intensity normalization, spatial filtering with a 6mm FWHM Gaussian kernel. We then denoised all preprocessed data for motion-related artefacts. We used ICA-AROMA to detect motion-related artefacts in single-subject data from ICA components extract by MELODIC and subsequently regressed these artefacts out of the data using fsl_regfilt (Beckmann and Smith, 2004; Pruim et al., 2015a, 2015b). Finally, we regressed out mean signals from CSF and white matter, and applied a 0.01Hz temporal high-pass filter (Gaussian-weighted least square straight line fit to the data).

For each participant, all acquisitions were registered to its high-resolution T1 image using Boundary-Based Registration (BBR) available in FSL FLIRT (Jenkinson and Smith, 2001; Jenkinson et al., 2002). All high-resolution T1 images were registered to MNI152 space using 12-dof linear registration available in FLIRT and further refined using non-linear registration available in FSL FNIRT (Andersson et al., 2007). Transformations were not applied. Instead we used the inverse of the obtained transformations to bring a hierarchical atlas of brain regions to the participant’s native space (see below).

### 2.3. Connectome atlas

For each functional imaging scan we defined connectivity matrices using regions defined in a hierarchical whole-brain functional atlas (van Oort et al., 2017). This atlas contains 185 non-overlapping regions and was defined through Instantaneous Connectivity Parcelation (ICP, (van Oort et al., 2017)) as applied to resting state fMRI data of 100 participants of the Human Connectome Project (HCP; (Glasser et al., 2016; Van Essen et al., 2012)). In short, ICP aims to parcel larger regions into subregions based on signal homogeneity, where the optimal number of subregions is determined based on split-half reproducibility at the cohort level.

Figure 1 illustrates the hierarchical brain atlas, where areas were grouped in 11 higher-level networks: 9 resting state networks (visual1, visual2, motor, right attention, left attention, auditory, default mode network (DMN), fronto-temporal and cingulum), and 2 networks based on anatomical structures, i.e., the subcortical areas, and the cerebellum. These higher-level networks respectively contained 19, 12, 22, 22, 18, 8, 18, 13, 7, 23, and 23 subregions, resulting in a total of 185 initial parcels.

**Figure 1.**
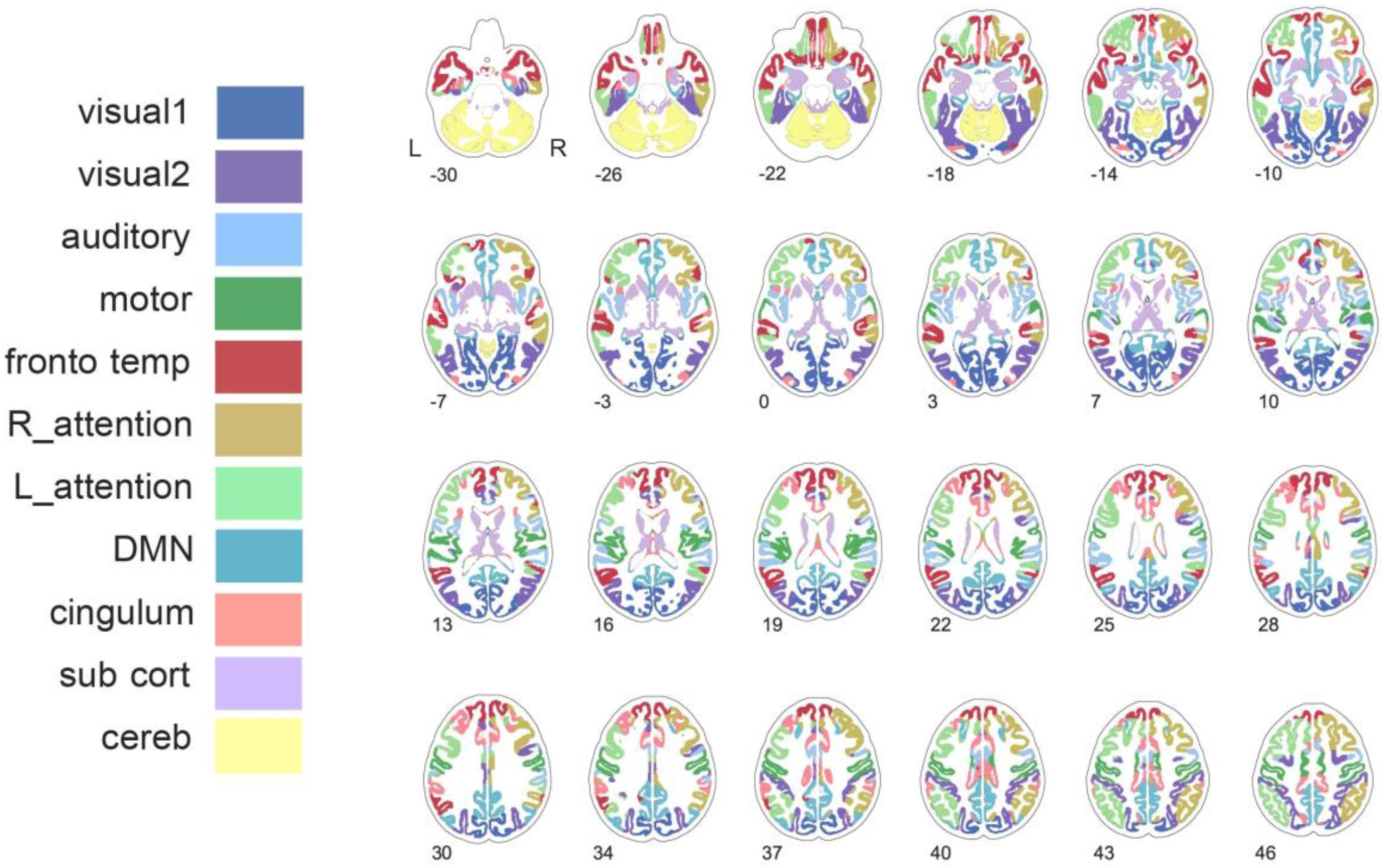
ICP atlas with 179 areas represented in their corresponding top-level networks. R_attention: right attention network; L_attention: left attention network; DMN: default mode network; sub cort: subcortical regions; cereb: cerebellum.

Connectivity matrices were calculated in each participant’s native space for each of the functional scans. To this end we transformed the atlas to each participant’s native space using the inverse of the anatomical to MNI152 non-linear warp, and the inverse of the linear transformation of the functional image to the participant’s high resolution anatomical image. Voxel-membership in brain parcels was established on the basis of majority overlap. Atlas areas that were on average across our population > 50% outside of the brain were rejected from further analyses. As a result, we used 179 areas, shown in Figure 1 colour coded by their associated top-level network, to compute connectivity matrices.

### 2.4. Connectivity calculation

For each participant and each task (rFMRI, WM, REWARD, STOP) we calculated 179x179 connectivity matrices, by cross-correlating the time series of all regions in the atlas. We obtained each region’s time series through multivariate spatial regression, using all 179 regions as regressors and each task’s preprocessed time series as dependent variable. The resulting regional time series were demeaned. Using these time series, we calculated 179x179 partial correlation matrices through inverting covariance matrices estimated by the Ledoit-Wolf normalization algorithm (Brier et al.; Ledoit and Wolf, 2004) as implemented in nilearn (<u>http://nilearn.github.io/</u>). Finally, all pair-wise correlations were Fisher *r*-to-*Z* transformed.

To allow comparing connectivity values between acquisitions and account for the potentially substantial differences in the temporal degrees-of freedom, we normalized the distribution of connectivity values within each connectivity matrix. We fit a mixture distribution under the assumption that the evidence for a non-zero connection is unrelated to the spatial location of the nodes and that non-zero connections are sparse. Further, we assume that there is a sufficient total number of nodes so that the distribution of values for non-existing edges in the network can be used to estimate the within-subject null distribution of non-existing connections. Specifically, we modelled the obtained connectivity values per task using a Gaussian-gamma mixture model (Beckmann et al., 2005; Feinberg et al., 2010)( Bielczyk 2017) and used the main Gaussian, i.e., the one fitting the body of the distribution, to normalize our connectivity values. In practice, we applied mixture modelling to each connectivity matrix and subsequently normalized the connectivity values by subtracting the mean and dividing by the standard deviation of the obtained Gaussian model. As a result, and despite differential loss in temporal degrees-of-freedom due to the partial correlation calculation, the values within the normalized, *Z*-transformed partial correlation matrices are readily comparable across participants and tasks.

Finally, to allow interpretation of the task-based connectivity matrices in terms of the deviation from the functional baseline defined as connectivity during the resting state, we further standardized each participant’s task-based connectivity matrix. Specifically, we standardized each individual-level pair-wise correlation by subtracting the corresponding individual pair-wise correlation obtained during rest for the same participant. As such, each task-based pair-wise correlation or edge quantifies how connectivity for that edge differed from that edge’s connectivity during the resting state. As a result, after standardization, we obtain for each participant an individual connectivity matrix for each of their task acquisitions. We refer to these matrices as *task potency matrices*, quantifying for each edge how strongly the task-based connectivity was modulated away from its resting state baseline. For each task, we finally create group-level task potency matrices by averaging across participant matrices and multiplying by the root mean square of the number of participants to avoid bias in between-task comparisons related to the number of observations in each task.

### 2.5. Task-based Fingerprints

To compare those connections that characterize a task’s functional fingerprint across different tasks we selected, for each task, those edges that showed a task potency that significantly deviated from rest. To this end, we converted the group-level task potency matrices to *Z*-statistic matrices by subtracting the mean and dividing by the standard deviation calculated for each task matrix. For each task, we selected those edges where |*Z*| >= 2.3. We refer to those edges as task-based fingerprints. The task-based fingerprints form the basis to define task sensitivity and task specificity of each edge in the fingerprint.

### 2.6. Network-based summary metrics

As described above, the hierarchical ICP atlas defines 179 areas as subdivisions of 11 large-scale networks. Accordingly, next to reporting at the level of individual areas, we can average across edges within each network to summarize potency, sensitivity, and specificity at the network level. We can differentiate edges that link areas within a network (within-network edges) from edges that link areas between two different networks (between-network edges). In order to compare between networks, we corrected for the number of edges averaged over the different 11x11 interactions by multiplying each average by the root mean square of the number of edges within or between two networks. By comparing the within-and between-network connections we assessed whether a task was associated with specific networks or resulted in an overall, diffuse modulation of connectivity. In practice, to derive network-level scores, we calculated the percentage of selected edges included in each network. This was done for each entry in the 11x11 network connectivity matrix, and allowed quantifying the selection of edges at the within-(diagonal matrix entries) and between-network (off-diagonal matrix entries) level.

**Figure 2.**
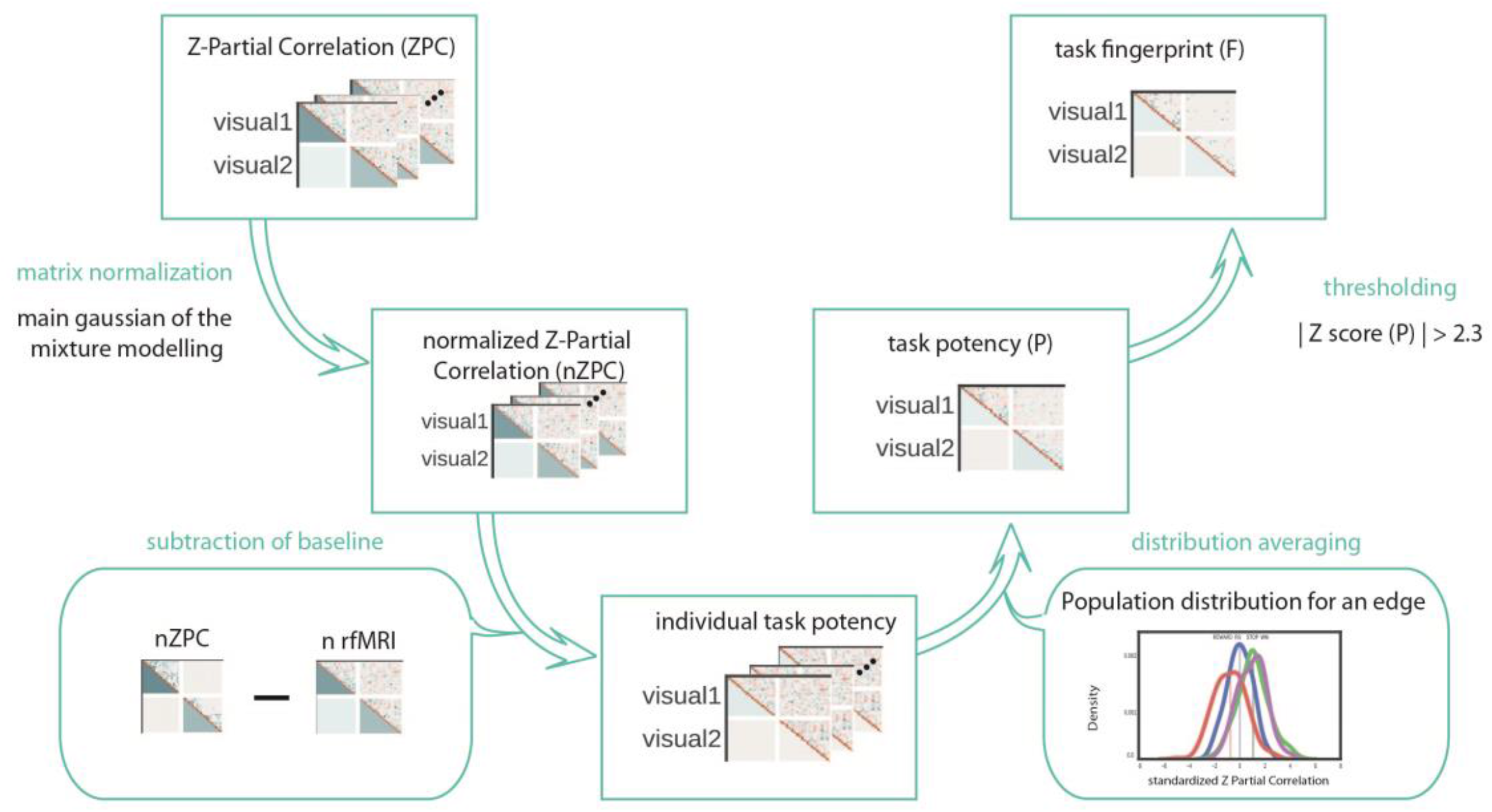
Analysis framework to obtain connectivity-based task fingerprints. The framework starts at the participant-level with obtaining a partial correlation matrix (Fisher-Z transformed), which is normalized, and subsequently standardized by that participant’s resting state connectivity (subtraction of baseline), resulting in individualized task potency matrices. A group task fingerprint can be obtained by averaging the individual task potency matrices and thresholding based on the z-score of the group potency.

### 2.7. Task Potency, Sensitivity, and Specificity

**Figure 3.**
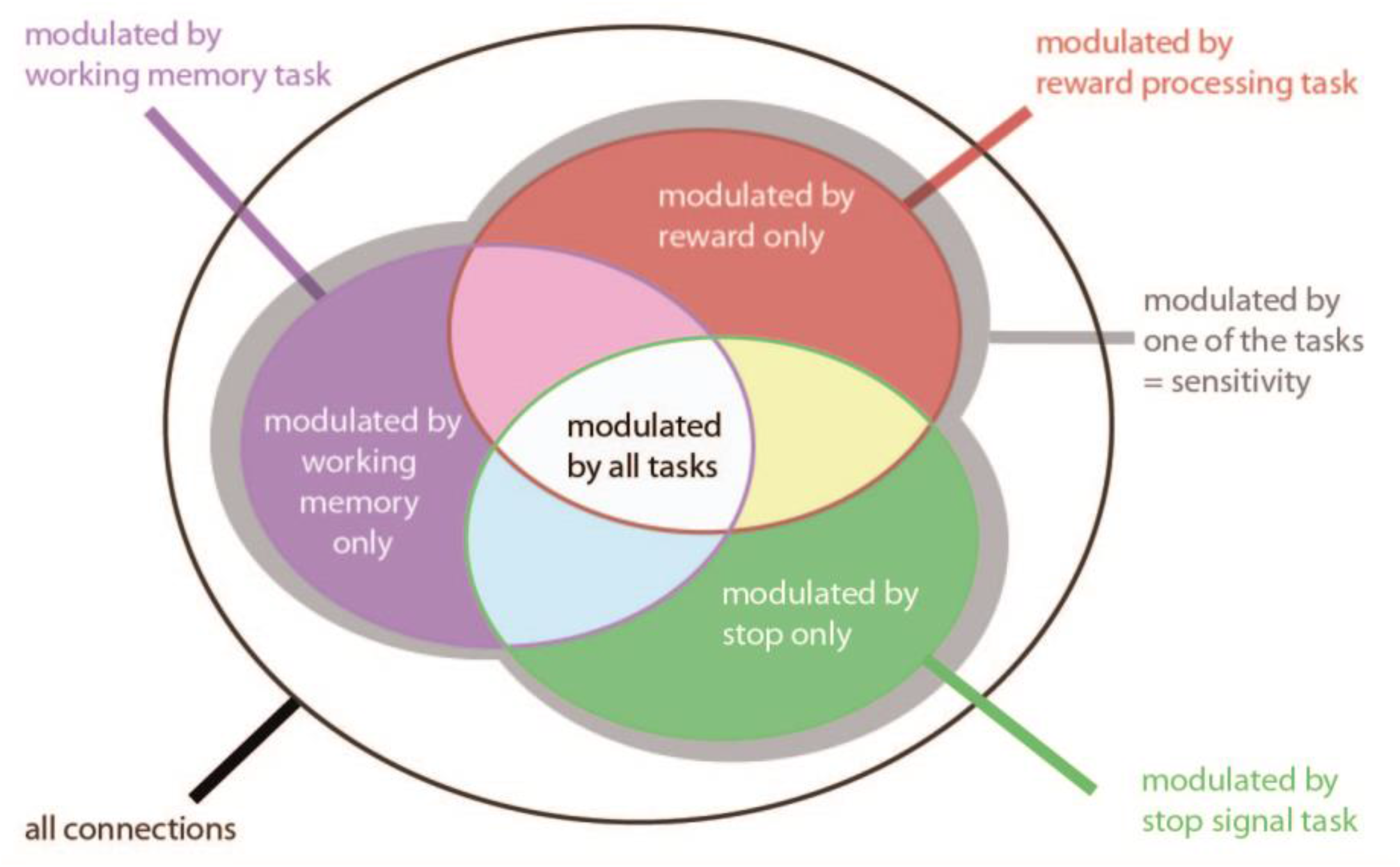
Overview of sensitivity and specificity as applicable to each edge within a task potency matrix.

Figure 3 illustrates how we can characterize each edge within the task potency matrix in terms of its sensitivity and specificity to the different tasks included in our investigation. An edge was regarded ***sensitive*** to task modulation when selected (i.e., a task potency above the |Z|>=2.3 threshold) in at least one of the tasks. An edge can be sensitive to modulation by several tasks, yet with a different level of potency in one task compared to another. However, this is not considered when assessing sensitivity.

Task ***specific*** edges were those edges that were selected for one task only. Note that specificity is determined by the collection of available tasks. Moreover, the bigger the difference in cognitive constructs between tasks, the easier it is to define specificity of involved edges.

Within the edges showing common sensitivity we further differentiate tasks based on their ***potency***, as an index of their ability to enlist a sensitive edge. To this end, we assessed whether one task was significantly more potent than others by comparing the distribution of task potency values for that edge across participants, between tasks. This comparison relies on the normalization of all matrices prior to standardization by the resting state connectivity (see supplementary Figure 2). We quantified the difference in potency between sensitive tasks by computing Cohen’s *d* between the obtained distributions. We required a minimum difference of 0.3 in Cohen’s *d* in order to decide which task specifically potentiated an edge. If all three tasks similarly potentiated an edge (i.e., the difference in Cohen’s *d* between them was <0.3) it was labelled as “common”. In contrast, if only two tasks similarly potentiated an edge, the edge was labelled as “undefined”, as it is neither specific to one task, nor common to all. The same method was used to label maximum task-related up- or down regulation in connectivity in the 11-network framework. In this case, the input were edges from the 11x11 matrices for each participant corresponding to the sum of potency across selected edges for each network. Finally, we investigated maximum task potency at the level of areas (i.e., columns in our connectivity matrices). To this end, we summed potency for group-level selected edges across each area’s 179 connections.

### 2.8. Reproducibility of the edge selection procedure

Every single participant has associated within-subject differences relative to the cohort-derived group connectivity matrix. In order to assess reproducibility of our group-level fingerprint pattern, we defined individual task fingerprints, applying the same selection procedure as above, but applied to the individual task potency matrices (i.e., select those edges with a |*Z*|>=2.3 in the individual task fingerprints). This enables us to quantify subject-specific variations in the edge selection and thereby permits quantification of reproducibility across participants. We indexed the number of times an edge was selected across participants. This proportion is interpreted as the reproducibility of an edge’s potency.

## 3. Results

### 3.1. Task-Based Fingerprints

Overall, the connectivity structure derived from each task was very similar to the connectivity structure derived from the resting state scan. Figure 4 illustrates this for the reward task (see supplement Figure 1 for the other tasks). To evaluate connectivity sensitivity to each task we created task-based fingerprints by standardizing the task connectivity by the resting state connectivity, resulting in a matrix quantifying each edge’s functional potency, as illustrated in Figure 4C. This task-based fingerprint served as the basis to identify sensitive edges and assess their task specificity.

**Figure 4.**
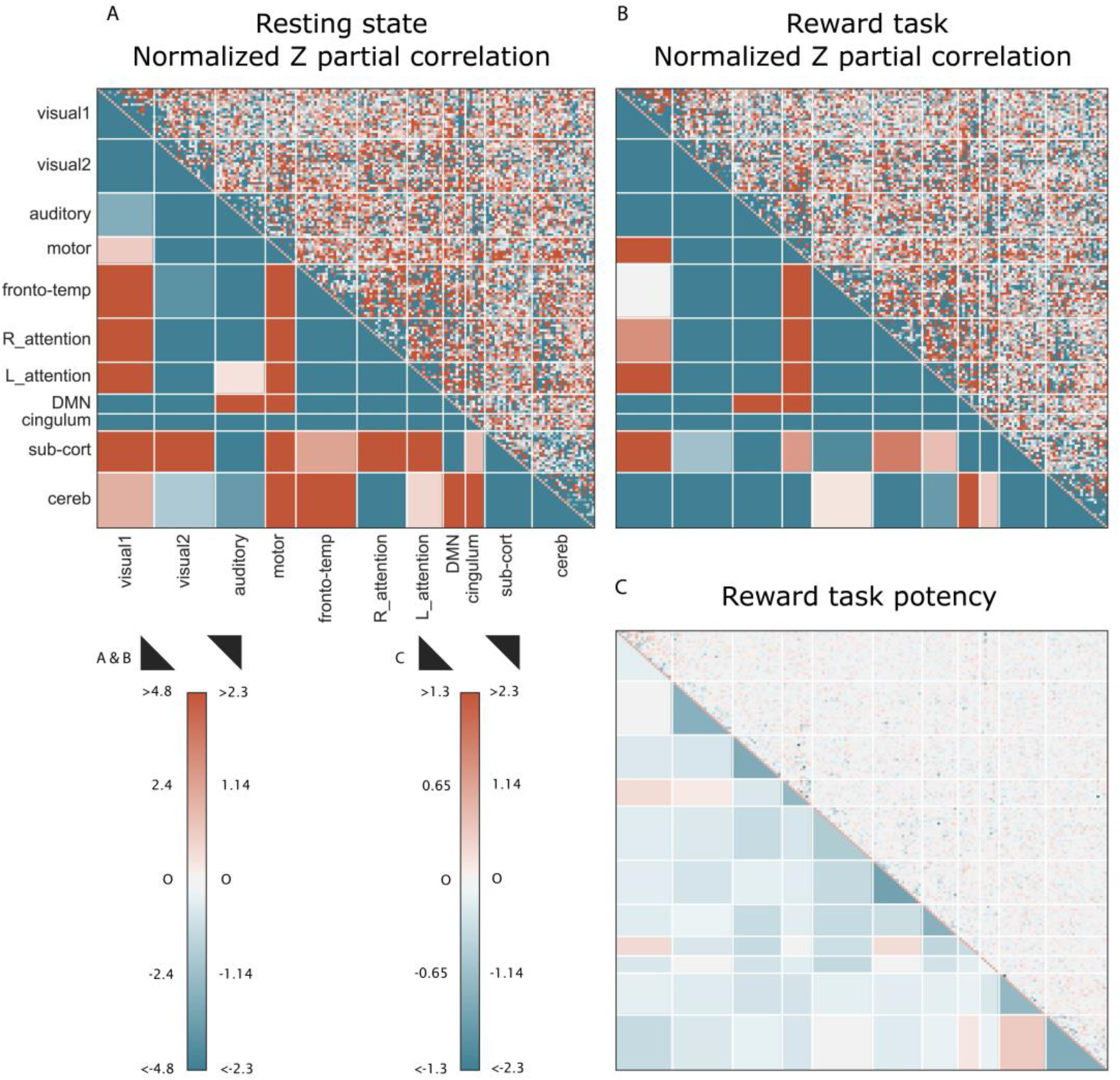
Illustration of connectivity matrix calculation for the reward task. **A**: Normalized resting state Z partial correlation averaged across the population; **B:** Normalized Z partial correlation averaged across the population; **C:** Average reward task potency across the population. Upper triangle displays the 179x179 connectivity fingerprints; lower triangle displays the average summary per network. R_attention: right attention network; L_attention: left attention network; DMN: default mode network; sub cort: subcortical regions; cereb: cerebellum. The normalized Z partial correlation and task potency matrices for the two other tasks are display in supplementary Figure 3.

### 3.2. Task Sensitivity

Using each task’s functional fingerprint, we determined which edges were sensitive to modulation by that task by selecting those edges that showed a |*Z*|-score above 2.3 in the average potency across participants. Figure 5 illustrates the characteristics of the selected edges across the three different tasks.

We observed a higher prevalence of sensitive edges for within-network connections compared to between-network connections. In particular, the visual1 and motor networks showed a high percentage of within-network sensitive edges. In contrast, the number of sensitive between-network connections was considerably lower, with on average only 1.5% of edges selected across the 11 networks versus 11.1% of within-network connections. As such, within-network connectivity is ~7x more strongly modulated than between-network connectivity. We highlight the result for the cingulum network which included twice as many sensitive between-network connections compared to the other networks, relative to no within-network potentiation. When comparing the percentage of sensitive edges between tasks we observed that overall all tasks yielded similar percentages of selected edges, with the exception of the within-network DMN connections, where STOP did not yield any sensitive within-network edges.

Overall, we observed greater sensitivity to task modulation in within-compared to between-network connections. This result suggests that during task performance there is increased change in communication within a network, rather than connecting regions across the brain. This result supports the hypothesis that the brain strongly segregates information at the level of individual networks, while more weakly integrating information between networks (Deco et al., 2015; Jirsa et al., 2014). However, as indicated before, going beyond overall statements requires assessing the task-specificity of each sensitive edge.

**Figure 5.**
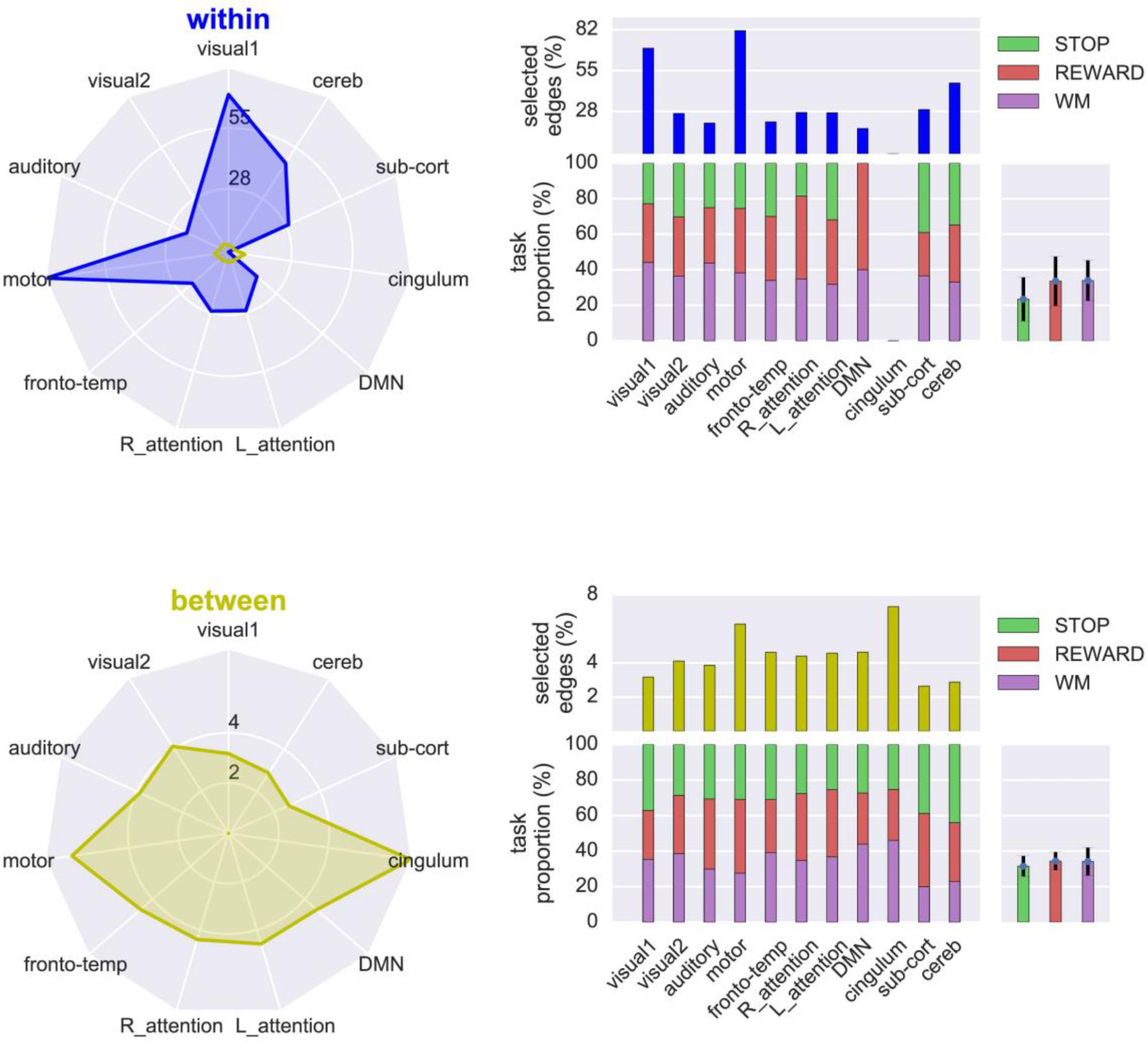
Radar plots of the percentage of edges showing connectivity changes (i.e., sensitivity) in each of the 11 networks. We observed a larger percentage of sensitivity for within-network connections (top), compared to between-network connections (bottom). As an example, 82% percentage of edges within the motor network exhibited sensitivity, compared to only 7% of its between-network connections. To allow direct comparison between both radar plots, we also show the between-network percentages on top of the within-network percentages in the top left plot. Bar plots on the right illustrate edge sensitivity for each task. For further detail on the percentage calculation we refer to supplementary Figure 4. R_attention: right attention network; L_attention: left attention network; DMN: default mode network; sub cort: subcortical regions; cereb: cerebellum.

### 3.3. Task Specificity

To disentangle overlapping connectivity modulation by the different tasks we define the task specific edges by dividing the obtained collection of sensitive edges into those that were modulated by one task only (i.e. specific to a certain task) versus those that were sensitive to modulation by several or all tasks (Figure 2). Figure 6 illustrates how the sensitive edges modulated by one task only or by all tasks were represented in the 11 networks. We observed that overall 62.2% of the sensitive edges were specific to a certain task, versus 17% that were modulated by all tasks. Note that this also means that 20.8% of sensitive edges was modulated by more than one, yet not all, tasks.

In the set of sensitive edges, we observed a difference in the level of specificity for within- vs between-network connections. Comparing the dark versus light coloured areas in the top row of Figure 6 it is evident that the ratio of specific versus common connections was smaller for the sensitive within-network connections (mean = 0.67 ± 0.45) compared to the ratio of specific versus common connections for the sensitive between-network connections (mean = 2.8 ± 2.94). As such, between-network connections are almost exclusively modulated in a specific fashion, where different tasks modulate different edges connecting networks to the rest of the brain. When taking into account the absolute percentages used to calculate the above ratio, we observed that the percentage of common connections was low for both the within-versus between-network connections (mean = 12.01 ± 8.01 vs mean = 6.35 ± 2.81). As such, the difference in the specific versus common ratio was driven by a difference in the absolute percentage of specific connections, with between-network connections exhibiting greater overall specificity. Note that the lower task-specificity of within-network connections implies a higher amount of connections that exhibited heterogeneous sensitivity to task.

We can further characterise the nature of the task-specific connections to assess how specificity is distributed across tasks and networks (Figure 6 bottom). While the between-network connections were more homogeneously distributed between tasks and across all networks, we observed greater variability in task specificity for the within-network connections, with some networks showing notable task-specificity. For REWARD, the attention networks and DMN showed higher specificity of within-network modulations. For STOP, the visual2, subcortical, and cerebellum edges exhibited most specific connectivity modulations. Finally, WM modulated specific connections within the visual1 and auditory networks. In contrast, we observed strongest task-specificity for between-network connectivity modulations involving the cingulum, which strongly preferred WM, while the between-network connections for the other networks did not exhibit a strong preference for a specific task.

In contrast to task-specific edges, about 16% of all task-sensitive edges were modulated by all three tasks (union between all tasks in Figure 2; dark areas in Figure 6, top row). Brain regions that showed the highest number of edges modulated by all tasks are represented in Figure 7. Apart from visual and motor regions where we expect shared modulation as all three tasks are presenting visual stimuli and request motor response, all tasks modulated edges involving regions that were part of the fronto-temporal and attention networks in our atlas. This modulation includes areas from inferior parietal lobe, bilateral frontal orbital cortex extending into Broca’s area, the temporal pole, amygdala, and enthorinal cortex. Interestingly, no regions from the DMN or the subcortical network were represented in the top selection of areas that potentiated edges across all tasks. Brain regions that showed the highest number of edges modulated by only one task are represented in supplementary Figure 5 (STOP), 6 (REWARD) and 7 (WM).

**Figure 6.**
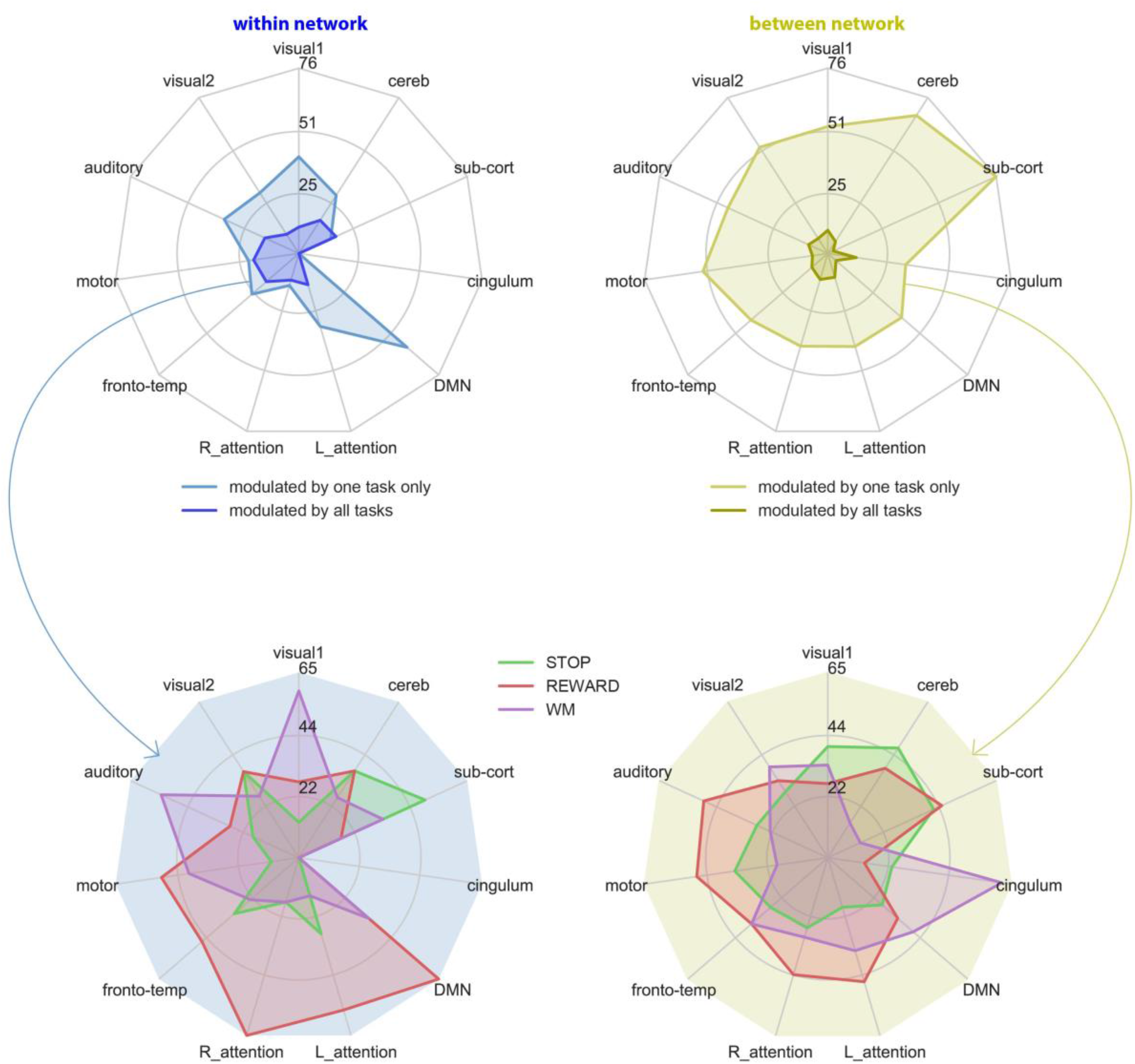
Percentage of sensitive edges for each task that end up to be specific to this task, e.g., percentage of edges selected in the group potency of a task that are not present in another task group potency selection. The percentage corresponding to the within- vs between-network connectivity is listed per network. Top row: overall results; bottom row: inflation of the edges modulated by one task only further differentiated per task. As an example: ~38% of the sensitive edges within the visual1 network were modulated by one task only. Of those 38% sensitive edges, about 55% was modulated only during WM performance, ~25% only during REWARD, and ~13% during STOP only. In contrast, ~10% of the sensitive edges within the visual1 network were modulated during performance of all tasks. Supplementary Figure 4 illustrates the reference edges in the percentage calculations. R_attention: right attention network; L_attention: left attention network; DMN: default mode network; sub cort: subcortical regions; cereb: cerebellum.

**Figure 7.**
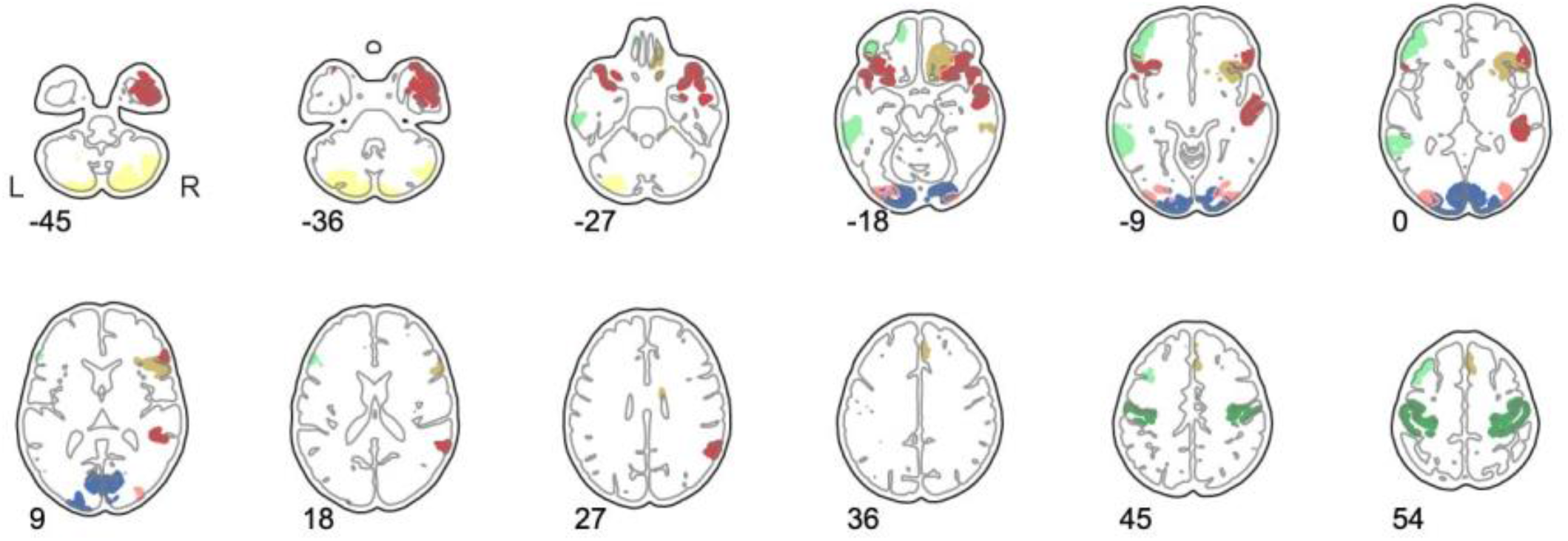
Brain regions with the highest number of edges commonly modulated by all three tasks. Here, we displayed the 10% brain regions with the largest sum of edges sensitive to all three tasks.

### 3.4. The most potent task

Even if each task’s set of specific modulations of connectivity reflects network enlistment by a specific experiment, this information is only accessible in comparison to other tasks. Additionally, to define edges as being task-specific brings limitations as soon as the number of tasks increases or when tasks share cognitive processes differentially involved in the experimental design. Therefore, we propose to move from the binary concept of specificity and sensitivity to a measure of connectivity potency to enlist a connection, a network, or an area of a task.

Using potency as a quantitative measure of the strength of enlistment of connections in a task, we can characterise which task potentiated an edge, a region, or a network most strongly (if at all). To this end, we assessed whether a task showed a significantly higher modulation of a given feature (i.e. an edge, a region, or a network) relative to other tasks by computing Cohen’s *d* between the population distributions of an edge’s task potency, corrected for the number of subjects included in each task. The task with the highest potency and a significant difference in Cohen’s *d* with other tasks was selected as the most potent task. We investigated the potency distribution of each edge separately (fig 8, upper triangle of the matrix). For networks (fig 8, lower triangle of the matrix) and areas (fig 8 brain slices), we looked at the sum of potency across selected edges within a network or involving a given brain region.

Potency of sensitive connections was mostly driven by a specific task, instead of showing common potentiation by all tasks. 66.7% of sensitive edges were more strongly modulated by one task only vs 14.0% of edges that were equally modulated by all tasks. In edges that were modulated by several tasks, 4.5% showed full task disassociation in that at least one of the modulating tasks exhibited a significant difference in their level of potentiation.

At the brain region and network-level, WM and REWARD offered most strong potentiation across the brain (Figure 8B). The reward task principally potentiated the right attention network as well as areas from the reward circuit (anterior cingulate cortex, prefrontal area, thalamus), whereas WM potentiated regions included in the DMN. The stop task mainly potentiated connections involving the visual 1 network and cerebellum.

**Figure 8.**
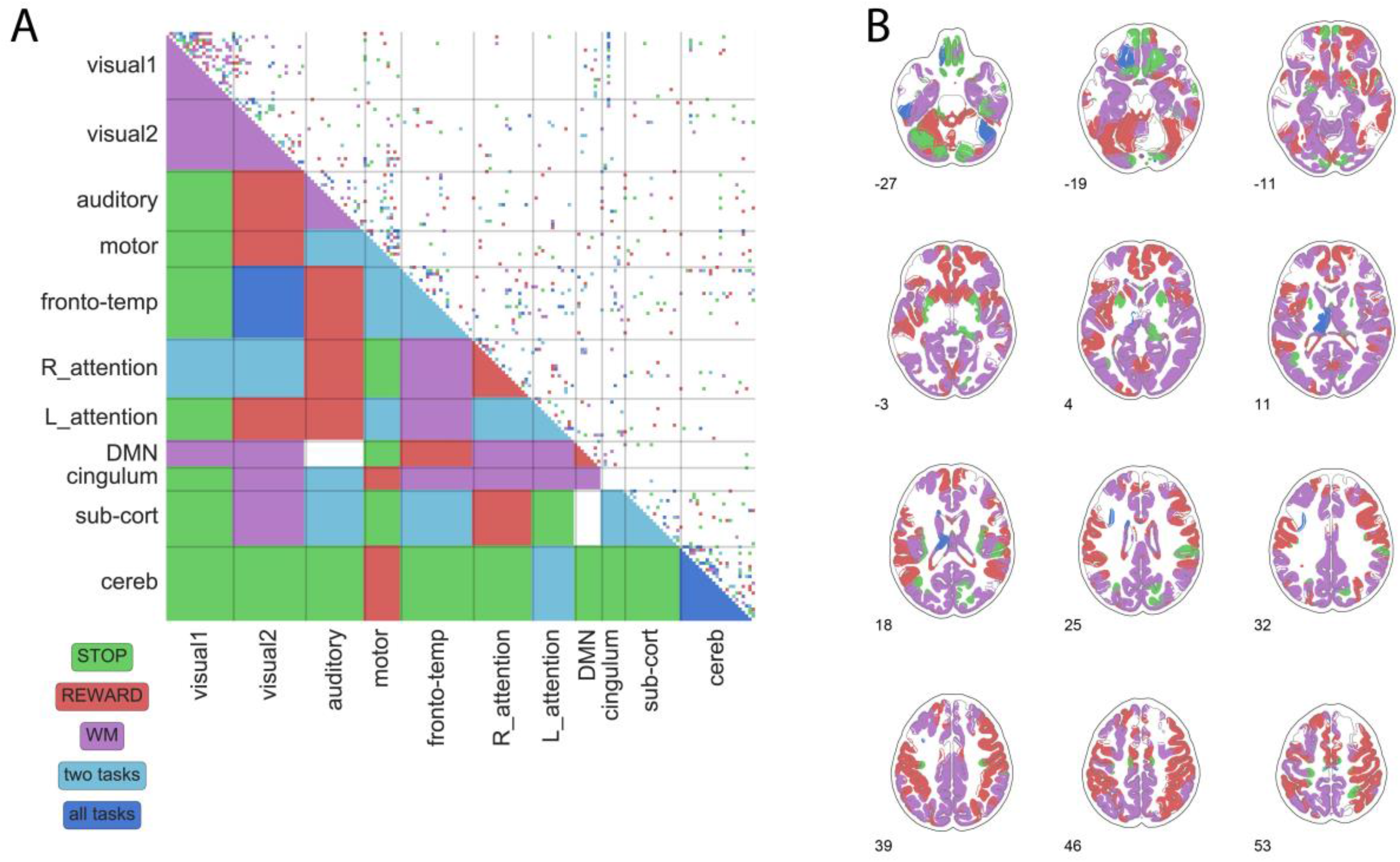
The most potent task illustrated for edges (A, upper triangle), networks (A (lower triangle), and brain regions (B). The brain slices on the right illustrate for each brain region which task on average potentiated edges involving this region the most. The same is shown for individual edges and at the network-level in the matrix on the left. Next to being most strongly potentiated by one specific task, we observed potency that was common to all tasks (i.e., no differentiation between tasks), or significant for some but not all tasks (two tasks). R_attention: right attention network; L_attention: left attention network; DMN: default mode network; sub cort: subcortical regions; cereb: cerebellum.

### 3.5. Reproducibility of the selection across individual fingerprints

The result in Figure 9 shows that different tasks exhibit a specific pattern of network potentiation, which can be accessed by comparing a set of different tasks. Nevertheless, a large proportion of sensitive edges are represented in all tasks. In order to establish the utility in evaluating individual task fingerprints in a reproducible manner, we studied the detection rate of edges sensitive to one or all tasks across task fingerprints obtained for individual participants. Specifically, we investigated whether the group selection was reproducible at the individual level and in particular how well the task-specific connections where represented at the level of individual participants. We defined the individual fingerprint by selecting edges that showed a |*Z*|-score higher than 2.3 in the individual-level task fingerprints. We computed the sum of selected edges across the population for each task. We observed a set of edges with high selectivity across participants for each task: 2.3% of edges within the union of individual masks were selected by minimum 15% of the population and in each of the three tasks. As shown in Figure 10, those edges mainly linked homotopic areas of each hemisphere, including bilateral motor areas, cerebellum, attention networks, visual1 areas, and bilateral putamen.

In contrast to the set of highly selected edges at the individual level, we note that the edge selection at the individual level showed substantial variability: 80% of sensitive edges are selected by less than 16% of the subjects (fig 9 dashed line). As indicated above, the most consistently selected edges between participants involved connections sensitive to all tasks. In contrast, highest inter-individual selection variability was found for task-specific edges as edges selected in only one task at the group level show a lower individual selection reproducibility than edges selected in all tasks at the group level (fig 10).

**Figure 9.**
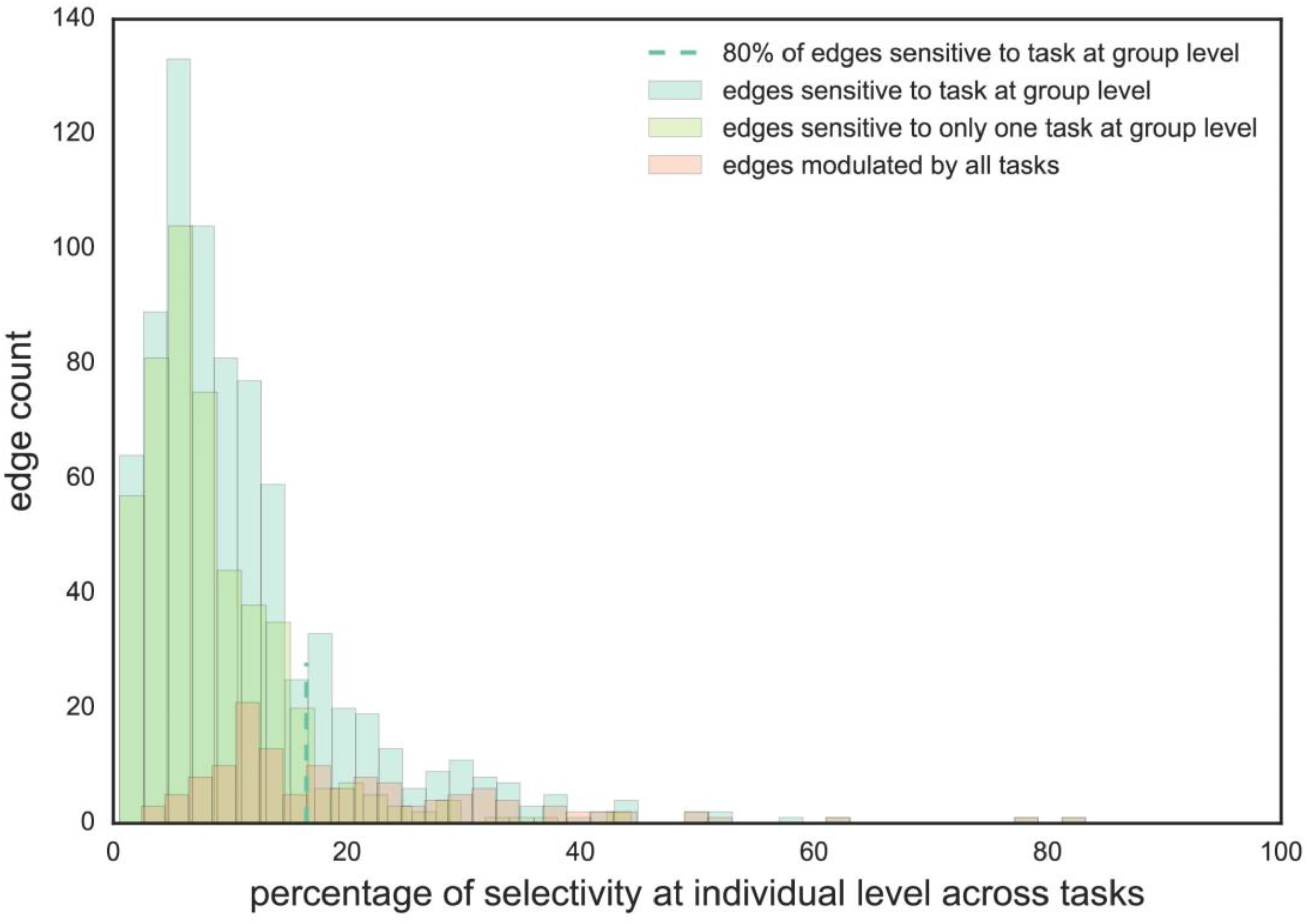
Distribution of selectivity of edges showing any sensitivity at the individual level. Distributions are shown for the selectivity of the corresponding edges at the group level for sensitivity to a task, sensitivity to one task only, and sensitivity to all tasks. The dashed line illustrates that 80% of the edges that were selected as sensitive to a task at the group level, were selected in about 18% of the individual task potency matrices. Through comparing the three distributions it is clear that overall edges modulated by all tasks were more consistently selected at the individual level.

**Figure 10.**
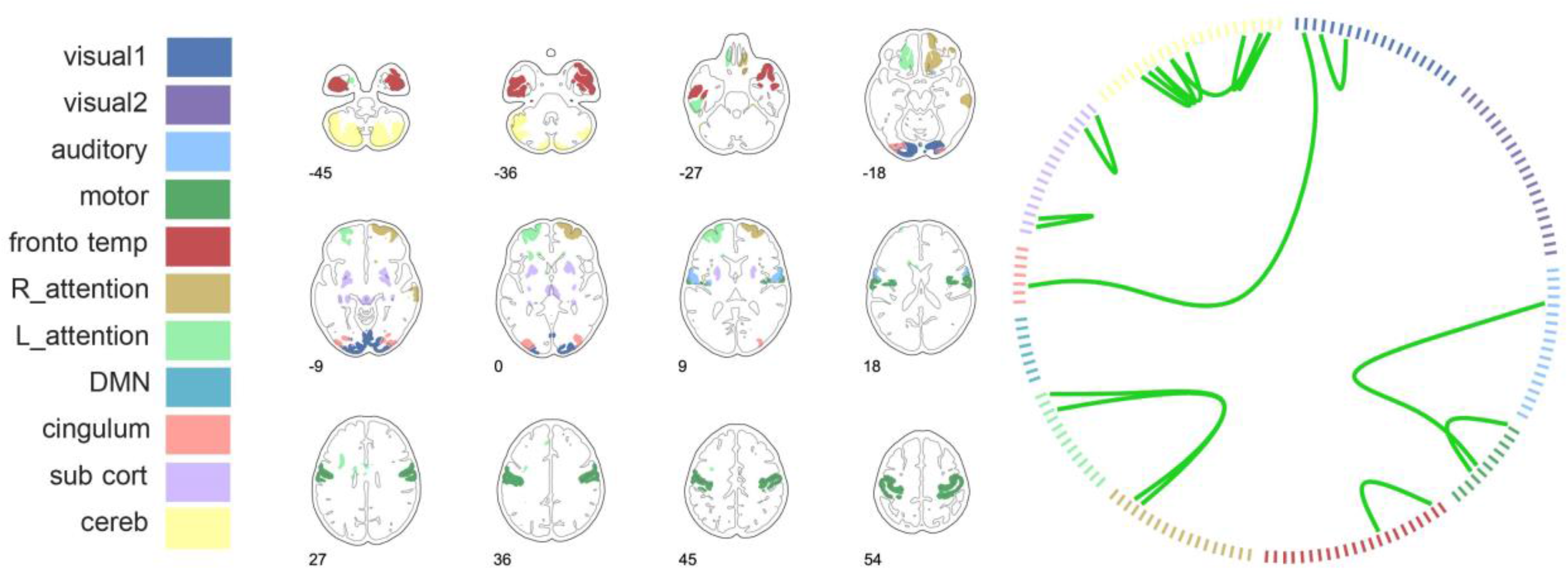
Edges selected for more than 20% of participants at the individual level. The circle displays significant connections that are being formed. The brain areas that these edges correspond to are represented in the axial slices. R_attention: right attention network; L_attention: left attention network; DMN: default mode network; sub cort: subcortical regions; cereb: cerebellum.

## 4. Discussion

When an individual engages in a task, the associated evoked activities build upon the brain’s ongoing activity, itself shaped by an underlying functional connectivity baseline (Fox et al., 2009; Smith et al., 2009; Tavor et al., 2016). Here, we show how this functional baseline architecture can be used to index task-dependent modulations, providing a means for quantitatively comparing evoked effects across different cognitive domains. This model incorporates the idea that functional connectivity observed under cognitive manipulation is task-specific with respect to its underlying resting state functional connectivity (Cole et al., 2014; Geerligs et al., 2015; Shirer et al., 2012; Smith et al., 2009). To facilitate understanding the building blocks of cognition we demonstrate that differential levels of localised sensitivity to task manipulation inform about the relative potency of a specific task.

Using this conceptual framework across multiple tasks, we show that task-induced changes in connectivity can be separated into common versus task-specific effects. We calculated task potency for three tasks (working memory task, response inhibition task, and reward processing task) in a large healthy population and showed that all tasks predominantly potentiated edges at the within-network level, i.e. more strongly connecting areas within networks, particularly those including lower-order sensory-motor regions. Such larger connectivity changes in primary sensory networks may highlight a more straightforward and automatic response to incoming stimuli, accompanied by standardized motor activity. This fits with the idea that visual and motor areas adhere to a highly constrained organization that is strongly evolutionary conserved, resulting in lower inter-individual variability (Mueller et al., 2013), but higher within-subject flexibility (Laumann et al., 2015).

Comparing tasks across multiple distinct cognitive domains allowed us to distinguish connections that were specific to each task versus those common to all manipulations. The task-specific connectivity modulations we observed were consistent with existing literature reporting on task-induced brain activity. For example, Figure 6 illustrates the high specificity of DMN connectivity in the working memory task, in accordance with the idea that DMN areas are involved in working memory (Piccoli et al., 2015; Pyka et al., 2009). By comparison, the stop signal task showed specific modulations involving areas typically observed in the inhibition networks (see also supplementary Figure 5; van Rooij et al., 2015), while reward processing specifically modulated putamen connectivity (see also supplementary Figure 6). Next to task-specific modulations, motor, visual, and higher-order cognitive regions including temporo-frontal areas showed sensitive yet unspecific involvement across multiple tasks (Figure 7). This result suggests that while these tasks probed different cognitive domains, they partially tap into similar cognitive processes resulting in similar connections exhibiting significant task potency.

Within the common connections, task potency amplitude can differ in accordance to the specifics of each task materialised into a continuum of potencies across different tasks per connection. Accordingly, we investigated different cognitive loads by computing the “most potent task” for selected connections. Overall, STOP seemed to be the least potent task in our study, yet supported by specific potentiation of visual and cerebellum connectivity with other networks. The two other tasks most potentiated brain regions and networks known to be involved in performance of those tasks: REWARD most potentiated the reward circuit (putamen, prefrontal cortex) and WM most potentiated the DMN. While we compared between tasks in the current study, a task potency continuum can also be obtained in relation to variation in cognitive load within a given tasks. Including such task designs would allow investigating the link between potentiation of connectivity and cognitive complexity.

Our task potency model is based on the idea that task activity builds on the brain’s inherent functional architecture as captured in large-scale resting state networks (Beckmann et al., 2005; Kelly et al., 2008; Mennes et al., 2010; Smith et al., 2009). Here, we showed preferential modulation of connections within those large-scale networks, while the limited number of modulated between-network connections exhibited greatest task specificity. This observation is consistent with the hypothesis that local processing is supported by out-of-network connections during task performance (Gratton et al., 2016). The idea that the resting functional architecture provides a common baseline further supports the need for a full and independent resting state acquisition to allow capturing the baseline functional landscape across the frequency range, as using in-task-OFF-block-resting-state data will be constrained by the task mind-set and relaxation of task potentiation.

To resolve and potentially quantify how specific a give task-activation pattern is for a cognitive function, current implementations to reverse inference typically rely on mining available literature (Hutzler, 2014; Poldrack, 2011; Schwartz et al., 2013; Varoquaux and Thirion, 2014). In contrast, our approach to compare each task against a resting baseline and to extract the amplitude of modulation as task potency at the cohort level enables comparing tasks using forward inference in a standardized space. This allows assessing connectivity specificity while avoiding potential literature bias or strong a priori models (study design, HRF response), albeit at the cost of being restricted to the typically smaller number of within-study cognitive domains being probed. We propose to integrate potency to define building blocks of cognition and integrate results from multiple task to solve how cognition emerges from the combination of these blocks. Comparing task potency amplitudes enabled us to define task-specificity that could in turn be used as a basis for a reverse inference framework.

While offering an interesting framework to study task fingerprints and connectivity specificity, we observed great variability in task potency across individuals. Task-related functional connectivity yields potential to understand individual variability in performance or task-related individual markers, as it has been successfully used to categorize tasks (Shirer et al., 2012), to predict performance (Cole et al., 2016), or to predict task-induced activity (Tavor et al., 2016). In our investigation we were unable to find strong associations linking task potency to task performance. Yet, supplementary Figure 8 (STOP), 9 (REWARD) and 10 (WM) illustrate that when correlating task potency and a corresponding task performance value that edges showing the strongest behaviour-potency correlations were in fact linked to task-relevant areas. Moreover, understanding task-related modulations might enable predicting functional connectivity in individuals that deviate from the norm, e.g., in a pathological response or using an alternative strategy to perform a task. Indeed, defining task potency relative to an individual resting state baseline is relevant for clinical applications where we cannot assume that individuals with pathologies have a similar baseline architecture as healthy control participants. As such, task potency might prove an interesting feature for cohort stratification, e.g. within the framework of normative modelling (Marquand et al.), aimed at characterizing how individual participants differs from a large normative range in regards to multiple brain-behaviour relationships.

In conclusion, our task potency framework quantifies task-induced connectivity changes relative to the resting-state baseline in order to index specificity of the brain’s functional architecture. Here, we showed that while general task performance relied mainly on within-network interactions, task specificity related to network interactions involving a close exchange between functional networks in both the cortex and subcortical structures. Using the potency framework we can address how function emerges in response to a task as well as how the brain’s baseline functional architecture influences cognitive operations. As such, the potency of our model lies in its ability to unfold the brain’s fluctuations in order to disentangle the common versus specific building blocks of cognition.

## Acknowledgments

This work was supported by a Marie Curie International Incoming Fellow-ship under the European Union’s Seventh Framework Programme (FP7/ 2007–2013), Grant No. 327340 (Brain Fingerprint) (to MM); The Netherlands Organization for Scientific Research (NWO) Grant No. NWO-Vidi 864-12-003 (to CFB); and Wellcome Trust UK Strategic Award [<u>098369/Z/12/Z</u>] (CFB). NWO Brain & Cognition: an Integrative Approach grant 433-09-242 (to JB), NWO National Initiative Brain & Cognition 056-13-015 (to JB), the EU FP7 grants TACTICS (278948) (to JB), IMAGEMEND (602450) (to JB), MATRICS (603016) (to JB) and AGGRESSOTYPE (602805) (to JB), and EU IMI grant EU-AIMS (115300)(to JB). The NeuroIMAGE project was supported by NIH Grant R01MH62873 (to Stephen V. Faraone), NWO Large Investment Grant 1750102007010 and ZonMW Grant 60-60600-97193 (to Jan Buitelaar), and grants from Radboud University Nijmegen Medical Center, University Medical Center Groningen and Accare, and VU University Amsterdam.

